# *Zymomonas mobilis* ZM4 utilizes an acetaldehyde dehydrogenase to produce acetate

**DOI:** 10.1101/2021.11.09.468001

**Authors:** Magdalena M. Felczak, Michaela A. TerAvest

## Abstract

*Zymomonas mobilis* is a promising bacterial host for biofuel production but further improvement has been hindered because some aspects of its metabolism remain poorly understood. For example, one of the main byproducts generated by *Z. mobilis* is acetate but the pathway for acetate production is unknown. Acetaldehyde oxidation has been proposed as the major source of acetate and an acetaldehyde dehydrogenase was previously isolated from *Z. mobilis via* activity guided fractionation, but the corresponding gene has never been identified. We determined that the locus ZMO1754 (also known as ZMO_RS07890) encodes an NADP^+^-dependent acetaldehyde dehydrogenase that is responsible for acetate production by *Z. mobilis*. Deletion of this gene from the chromosome resulted in a growth defect in oxic conditions, suggesting that acetaldehyde detoxification is an important role of acetaldehyde dehydrogenase. The deletion strain also exhibited a near complete abolition of acetate production, both in typical laboratory conditions and during lignocellulosic hydrolysate fermentation. Our results show that ZMO1754 encodes the major acetaldehyde dehydrogenase in *Z. mobilis* and we therefore rename the gene *aldB* based on functional similarity to the *Escherichia coli* acetaldehyde dehydrogenase.

**Importance:** Biofuel production from non-food crops is an important strategy for reducing carbon emissions from the transportation industry but it has not yet become commercially viable. An important avenue to improve biofuel production is to enhance the characteristics of fermentation organisms by genetic engineering. To make genetic modifications successful, we must gain sufficient information about the genome and metabolism of the organism to enable rational design and engineering. Here, we improved understanding of *Zymomonas mobilis*, a promising biofuel producing bacterium, by identifying a metabolic pathway and associated gene that lead to byproduct formation. This information may be used in the future for genetic engineering to reduce byproduct formation during biofuel production.

## Introduction

*Zymomonas mobilis* is an α-proteobacterium that produces ethanol from glucose with a very high specificity and may be useful for biofuel production (1). Under optimal conditions, *Z. mobilis* converts glucose to ethanol at up to 97% of the theoretical yield, converting only a small fraction of the carbon into byproducts or biomass (2, 3). However, under less favorable conditions, *Z. mobilis* generates byproducts, including acetaldehyde and acetate, especially when exposed to oxygen (4–6). Stressors present in lignocellulosic hydrolysates may also lead to byproduct formation. We previously observed that *Z. mobilis* produces 5-10 mM acetate when grown in anaerobic lignocellulosic hydrolysate (7). Acetate production during lignocellulose fermentation is problematic because it reduces the efficiency of biofuel production and further increases the acetate concentration. Acetate is formed by many biomass hydrolysis methods, leading to high acetate concentrations in lignocellulosic hydrolysates, which are often inhibitory to the fermentation organism (8, 9). Because of the already inhibitory concentrations and low value of acetate, generation of this molecule during hydrolysate fermentations is highly undesirable.

Although acetate production by *Z. mobilis* has been well-documented, the metabolic pathway that leads to significant acetate secretion is unknown (10). One possible source of acetate secreted by *Z. mobilis* is oxidation of acetaldehyde, which is an intermediate in the ethanol production pathway. Acetaldehyde is typically undetectable in anoxic cultures of *Z. mobilis* but accumulates in oxic cultures because of a redox imbalance. NAD(P)H that is oxidized by the respiratory chain is not available to reduce acetaldehyde to ethanol, leading to a buildup of acetaldehyde (**Figure 1**). Under oxic conditions, oxidation of acetaldehyde to acetate provides a possible route for detoxification, with additional NAD(P)H being recycled by the respiratory chain.

**Figure 1.**
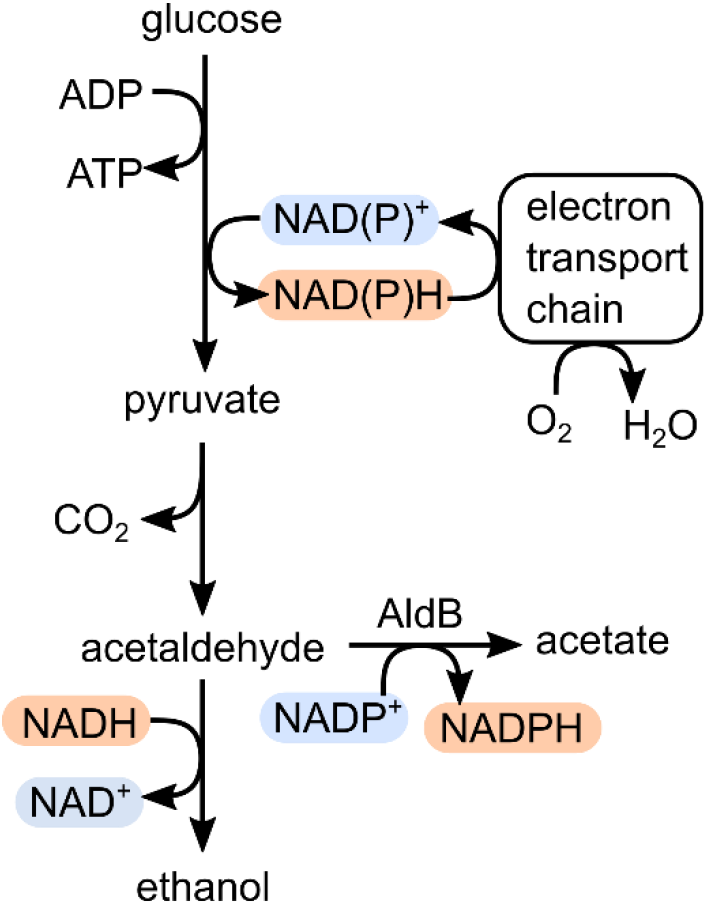
Central metabolism of *Z. mobilis,* including the acetaldehyde dehydrogenase encoded by *aldB*. Oxidized redox cofactors are highlighted in blue and reduced redox cofactors are highlighted in red to indicate the connections formed between oxygen and ethanol, acetaldehyde, and acetate formation.

An NADP^+^-dependent acetaldehyde dehydrogenase was discovered in *Z. mobilis* strain Z6 in 1989 *via* activity guided fractionation, suggesting that acetaldehyde oxidation could be a major source of acetate (11). However, there is no acetaldehyde dehydrogenase annotated in the *Z. mobilis* Z6 genome (12). Further, we have not found an annotated acetaldehyde dehydrogenase in any other *Z. mobilis* genome (7). As a result, most genome scale metabolic models for *Z. mobilis* only include acetate as a byproduct of less central reactions, such as amino acid biosynthesis (10, 13, 14). Only one model includes acetaldehyde dehydrogenase, although no gene evidence was provided for the reaction (15). Metabolic models of *Z. mobilis* tend to underestimate acetate production compared with experimental results (10), suggesting that a major source of acetate is missing. To address this gap in our knowledge of *Z. mobilis* metabolism, we identified a locus in the *Z. mobilis* ZM4 genome encoding an acetaldehyde dehydrogenase and determined that it is the main source of acetate production under laboratory growth conditions and during lignocellulosic hydrolysate fermentation. Here, we refer to the locus ZMO1754 as *aldB* due to the functional similarity to the *E. coli* gene of the same name. The ZM4 strain was used because it is the most promising strain for hydrolysate fermentation (7) and recently developed genome modification tools facilitated generation of a markerless deletion strain in this background (16).

## Materials and Methods

### Media and Chemicals

*Escherichia coli* strains were grown in Luria-Bertani medium (Miller, Accumedia). *Zymomonas mobilis* ZM4 was grown in ZRMG (1% yeast extract, 2% glucose, and 15 mM KH_2_PO_4_) or ZMMG (1.0 g KH_2_PO_4_, 1.0 g K_2_HPO_4_, 0.5 g NaCl, 1.0 g (NH_4_)2SO_4_, 0.2 g MgSO_4_ × 7 H_2_O, 0.025 g Na_2_MoO_4_ × 2 H_2_O, 0.025 g FeSO_4_ × 7 H_2_O, 0.010 g CaCl2 × 2 H_2_O, 0.001 g calcium pantothenate, 20.0 g D-glucose per 1 liter). Solid media contained 1.5% agar (Difco). Chloramphenicol was added to 100 μg/ml or 35 μg/ml, and spectinomycin to 100 μg/ml or 50 μg/ml for *Z. mobilis* and *E. coli*, respectively. Ampicillin was used at 100 μg/ml for *E. coli.* 2,6-Diaminopimelic acid (DAP, Aldrich) was added to a final concentration of 0.3 mM when needed. B-PER Reagent was from Thermo Scientific, Profinity immobilized metal affinity chromatography (IMAC) Ni-charged Resin was from Bio-Rad; all IMAC buffers contained 20 mM Tris HCl pH 8, 0.3 M NaCl, 10% glycerol and imidazole at 5 mM, 20 mM or 500 mM for IMAC A, B and C, respectively. Complete EDTA-free Protease Inhibitor tablets were from Sigma-Aldrich/Roche. Bradford Protein Assay Reagents 1 and 2 and Precision Plus Protein All Blue Prestained Protein Standard were from Bio-Rad. The anti His-tag mouse antibody was from GenScript (cat no. A00186). Horseradish peroxidase-conjugated rabbit anti-mouse antibody was from Sigma-Aldrich. TBST buffer is 20 mM Tris-HCl pH 7.5, 150 mM NaCl, 0.01% Tween-20. Clarity Western ECL was from BioRad. NADP^+^ sodium salt was from Sigma-Aldrich. NAD^+^ was from Amresco. Acetaldehyde, propionaldehyde, butyraldehyde, valeraldehyde, formaldehyde and glyceraldehyde were from Sigma-Aldrich/Millipore. Restriction enzymes, T4 ligase, Q5DNA Polymerase and HiFi DNA Assembly Master Mix were from New England Biolabs. Egg lysozyme was from Sigma-Aldrich and IPTG was from Goldbio. Oligonucleotides were synthesized by Integrated DNA Technologies (IDT).

### Bacterial strains and plasmids

*Zymomonas mobilis* ATCC 31821 (ZM4) was obtained from Dr. Robert Landick (University of Wisconsin, Madison). *E. coli* strains Mach 1 and BL21(DE3) pLysS, were from Invitrogen. *E. coli* WM6026 (*lacIq rrnB3* ΔlacZ4787 *hsdR*514Δ*araBAD*567 Δ*rhaBAD568* rph-1 attλ::pAE12(ΔoriR6K-cat:: Frt5), ΔendA::Frt uidA(ΔMluI)::pir attHK::pJK1006(ΔoriR6K-cat::Frt5; trfA::Frt) ΔdapA::Frt, (17)) was obtained from Dr. Patricia Kiley (University of Wisconsin, Madison). Strain ZM4 Δ*aldB* was constructed using the method described in Lal et al. (16). Briefly, 500-bp fragments directly upstream and downstream of *aldB* were PCR amplified with Q5 DNA Polymerase using the following pairs of primers: “ZMO1754 up” for upstream and “ZMO1754 dn” for downstream fragments (Table 2). The fragments were cloned into the SpeI restriction site of a non-replicating plasmid pPK15534 (16), digested with SpeI, using NEB Builder HiFi DNA Assembly Master Mix to get pPKΔZMO1754 (Fig S1). The insertion was confirmed by PCR. pPKΔZMO1754 was introduced into ZM4 by conjugation from WM6026/ pPKΔZMO1754, and subsequent selection for chloramphenicol resistance and DAP independence, as described in Lal et al. (16). Integration of pPKΔZMO1754 into the chromosome by homologous recombination in chloramphenicol resistant cells was confirmed by colony PCR. One colony with inserted plasmid was grown for about 10 generations in liquid ZRMG medium without chloramphenicol and 10,000 CFU were plated from an appropriate dilution onto ZRMG plates to get around 100 colonies per plate. Plates were incubated for at least 48 hours in aerobic conditions at 30°C and colonies were screened for loss of green fluorescence under a Blue Light Illuminator. Non-fluorescent colonies were checked for loss of *aldB* by PCR using primers *aldB* up F and *aldB* dn R (Table 2), followed by Sanger sequencing.

Plasmids and oligonucleotides are listed in Table 1 and Table 2, respectively. pET16b*aldB* was constructed by amplification of *aldB* from ZM4 genomic DNA using primers MF_17 and MF_18, and Gibson assembly into the pET16b amplified from primers MF_19 and MF_20. This assembly places *aldB* between BamHI and NdeI of pET16b under control of the T7 promoter and downstream of a 10-histidine tag (Figure S2). After transformation into Mach 1, plasmid was isolated from ampicillin resistant colonies and the sequence was confirmed by Sanger sequencing. Next, pET16b*aldB* was transformed to BL21(DE3) pLysS and IPTG-induced overexpression of His-tagged AldB was confirmed by Western blotting as described in His-AldB purification section. pRL*aldB* was constructed by amplification of *aldB* from ZM4 using primers MF_9 and MF_10 and assembled with pRL814 amplified with primers MF_11 and MF_12 using NEB HiFi DNA Assembly Mastermix. This assembly replaces the GFP gene in pRL814 with *aldB*. pRLHis-*aldB* was constructed by amplification of His-*aldB* from pET16b*aldB* using primers MF_23 and MF_24 and Gibson assembly with the pRL814 backbone amplified with primers MF_21 and MF_22. After transformation into Mach 1, isolated plasmids were sequenced by the Sanger method.

**Table 1.**
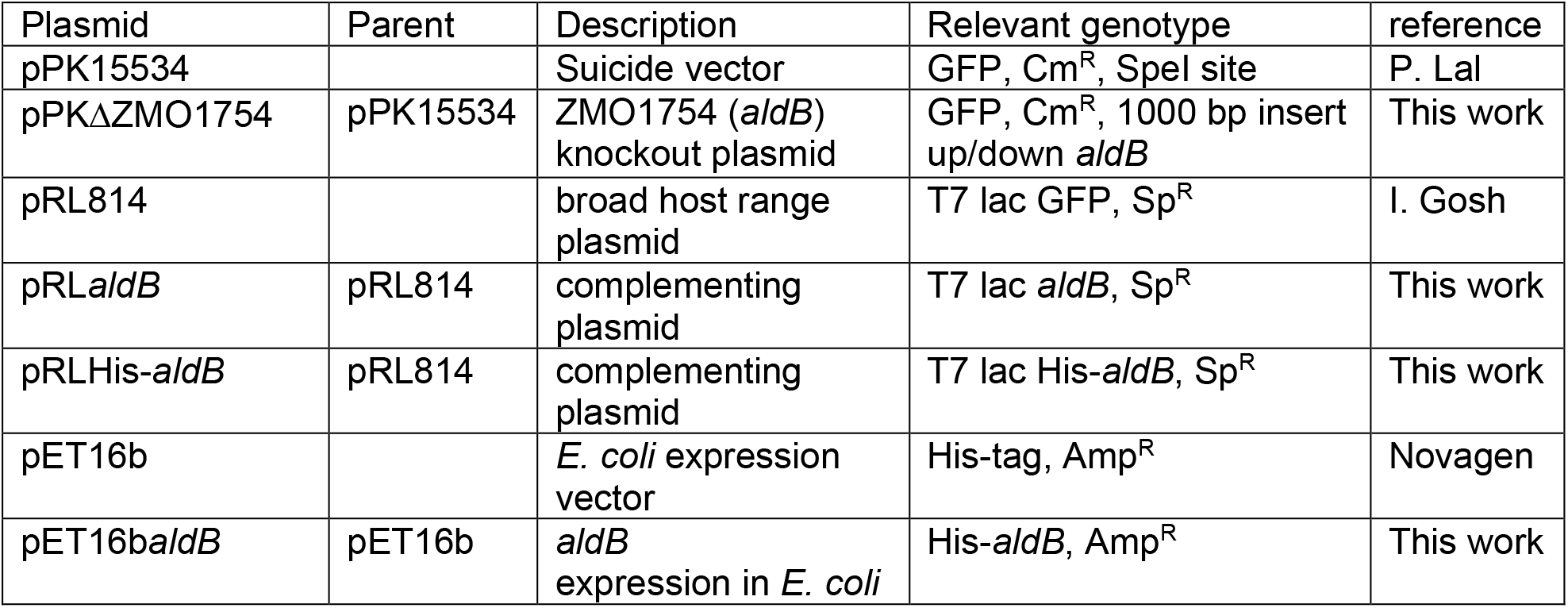
Plasmids.

**Table 2.**
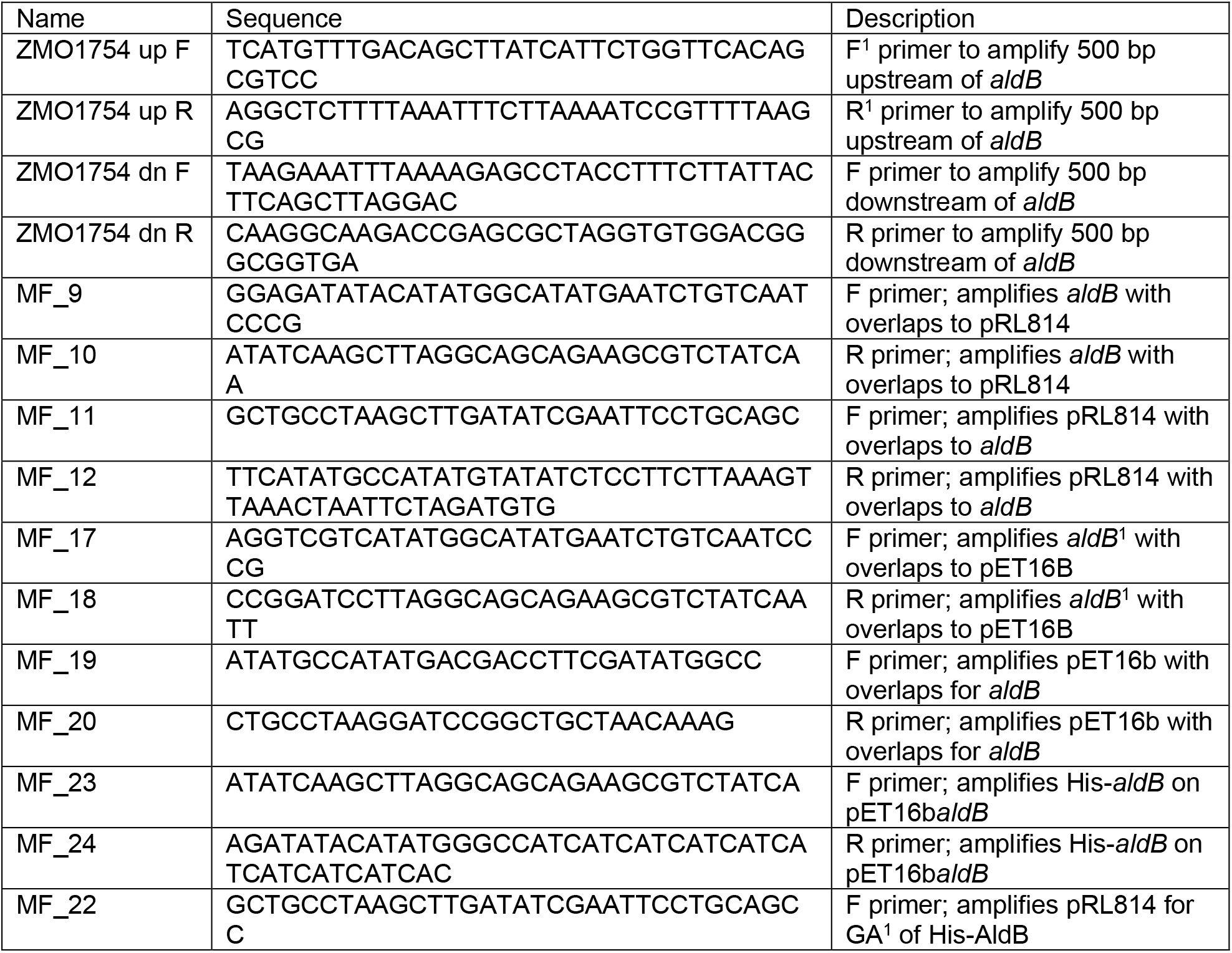
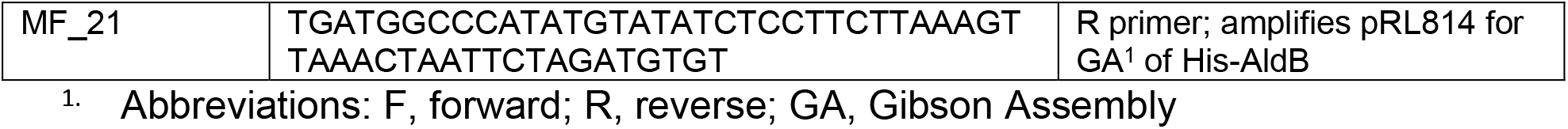
Oligonucleotides.

### Bacterial growth

ZM4 or Δ*aldB* were grown in ZRMG or ZMMG at 30°C, with shaking at 250 rpm. For anaerobic conditions bacteria were grown in Hungate tubes: 5 ml of media stored in an anaerobic chamber (Coy Lab Products) were inoculated from single colonies in the chamber and the tubes were closed with butyl-rubber stoppers and secured with aluminum crimps before removing from the chamber. For aerobic growth Hungate tubes were loosely covered with aluminum foil. Samples were removed periodically for OD_600_ measurements and HPLC analysis. Spectinomycin was added to cultures bearing pRL814, pRL*aldB* or pRLHis-*aldB* and IPTG was added as indicated.

### HPLC analysis

Samples were prepared by centrifugation at 14,000 rpm for 10 min in microfuge tubes at 4°C to remove cells and supernatants were transferred into 2 ml glass vials. Analysis was performed on Shimadzu 20A HPLC using Aminex HPX-87H (BioRad) column at 50°C. Separation of compounds was with 5 mM sulfuric acid at flow rate of 0.6 ml/minute. Compounds were identified by refractive index and concentrations determined from standard curves for external standards run in the same batch.

### *Soluble protein fraction from* Z. mobilis

50 ml ZRMG in 250-ml flasks were inoculated from fresh colonies of ZM4 or Δ*aldB* and grown aerobically with shaking at 250 rpm at 30°C to OD_600_ 1.0. Cells were harvested by centrifugation at 4°C at 8,000 rpm for 10 minutes (Sorval ST 8R, 75005709 rotor). Pellets were washed twice with 2 ml ice-cold 0.1 M Tris HCl pH 8.0 and resuspended in 0.25 ml of the same buffer. Lysis components were added to final concentrations: 20 mM EDTA, 1 mg/ml lysozyme, 4 mM DTT and Protease Inhibitor as recommended by manufacturer. Lysis was performed at 37°C for 20 minutes. Samples were spun in microfuge tubes at 17,000 rpm (Sorval ST 8R 75005715 rotor), for 30 minutes at 4°C. Clear supernatant (soluble protein fraction, FI) was gently transferred to new microfuge tubes and flash freezed in liquid nitrogen. FI was stored at −80°C.

### Purification of His-AldB by Ni-affinity chromatography

100 ml of LB containing ampicillin was inoculated from a single colony of *E. coli* BL21(DE3) pLysS bearing pET16b*aldB* and grown at 30°C with shaking at 250 rpm overnight. An overnight culture was diluted to OD_600_ = 0.1 in two liters of LB media with ampicillin in a 6-L flask and grown with shaking at 30°C to OD_600_ = 0.6. At this point, IPTG was added to a final concentration of 0.1 mM and growth was continued for two hours to OD_600_ = 2.5. Cells were harvested by centrifugation at 6,000 rpm for 10 minutes (GC-3 Sorval) at 4°C. Pellets were resuspended gently in 20 ml of ice-cold B-PER Reagent containing two tablets of Complete Protease Inhibitor and flash freeze in liquid nitrogen. On the next day, cells were thawed on ice and solid NaCl was added to a final concentration of 0.3 M and 100 mM imidazole to 5 mM. At this point cells were completely lysed. The soluble fraction was separated from membranes and cell debris by ultracentrifugation (Beckman, Ti 45) at 22,000 rpm for 40 minutes. Clear supernatant, containing soluble protein fraction (FI), was analyzed by SDS PAGE for the presence of His-AldB and the total protein concentration was determined using a Bradford assay. Activity of His-AldB was measured by an acetaldehyde dehydrogenase assay as described below. 11 ml of FI containing 14 mg/ml of total protein was mixed with 2.5 ml of Ni-charged IMAC resin equilibrated with IMAC buffer A and incubated on a rocker for 1 hour in 4°C. The entire slurry was loaded onto a 10-ml ThermoFisher disposable column under gravity flow at 4°C. The column was washed with 10 column volumes (CV) of IMAC A and 10 CV of IMAC B under gravity flow. His-AldB was eluted with 5 CV of IMAC C; 0.25-ml fractions were collected. Protein concentration in fractions was determined by the Bradford method and the purity of His-AldB was estimated by SDS PAGE followed by Coomassie staining and Western blotting with anti-His antibody. Dithiotreitol (DTT) was added to a final concentration of 10 mM to the peak fractions before freezing in liquid nitrogen. Fractions were stored at −80°C.

### SDS PAGE and Western blotting

The indicated amounts of Ni-chromatography fraction of His-*aldB* or soluble protein fraction was added to gel loading buffer (0.4 M Tris base, 30% glycerol, 1% SDS, 0.5% bromophenol Blue), containing 10 mM DTT to get final volume of 10 μl and boiled for 5 minutes. To get whole cell lysate, a pellet from 1 ml of cells at OD_600_ = 1.0 was resuspended in 0.1 ml of loading buffer and boiled for 10 minutes. Samples were loaded on 4–20% gradient Mini-PROTEAN TGX stain free gels (BioRad) and run in Tris/Glycine/SDS running buffer (BioRad). Gels were stained with Coomassie Brilliant Blue or proteins were transferred into nitrocellulose membranes in Trans-Blot-Turbo Transfer Buffer (BioRad) for 7 min in a Trans-Blot Turbo Transfer System (BioRad). Membranes were blocked with 3% BSA in TBST and incubated with mouse anti-His monoclonal antibody overnight in a cold room. Membranes were washed in TBST buffer for two hours and incubated with horseradish peroxidase-conjugated rabbit anti-mouse antibody for two hours. The membranes were washed and chemiluminescent detection was performed with Clarity Western ECL (BioRad) for 5 min.

### Standard acetaldehyde dehydrogenase assay

Reactions were performed in vitro in 1 ml of reaction buffer in 24-well plate using a Synergy H1 BioTek plate reader. The standard reaction buffer contained 50 mM Tris pH 8.0, 10 mM DTT, 22 mM acetaldehyde, 2 mM NADP^+^ or NAD^+^ and 50 mU of purified His-AldB or indicated volumes of soluble protein fraction from *Z. mobilis* or *E. coli* expressing *aldB* from plasmid. The reaction was performed at 30°C for 30 minutes. Acetaldehyde dehydrogenase activity was measured as reduction of NAD^+^/NADP^+^ to NADH/NADPH by monitoring absorbance at 340 nm.

## Results

To determine which gene in the *Z. mobilis* ZM4 genome encodes an ortholog of the previously purified acetaldehyde dehydrogenase from Z6, we searched for a locus with the following characteristics: i) the gene should encode a protein with an approximate size of 55 kDa, ii) the gene should be induced under aerobic conditions (11). Based on a recent multi-omic analysis of oxygen exposure in *Z. mobilis* ZM4 by Martien et al. (18), we hypothesized that locus ZMO1754 (also known as ZMO_RS07890) encodes the previously discovered acetaldehyde dehydrogenase. ZMO1754 encodes a protein with an expected size of 49.6 kDa, and both its transcript and protein abundance increased upon oxygen exposure (18). ZMO1754 is annotated as an NAD^+^-dependent succinate-semialdehyde dehydrogenase, suggesting a role as a redox enzyme. Further, an ortholog of ZMO1754 is present in the Z6 genome, indicating that it could encode the previously identified acetaldehyde dehydrogenase. The results below show that the protein encoded by ZMO1754 shares strong functional similarity to the *E. coli* gene *aldB*, which is also an NADP^+^-dependent acetaldehyde dehydrogenase (19). Due to the functional similarity, we propose to rename the locus ZMO1754 to *aldB* in *Z. mobilis* and refer to the gene as *aldB* throughout the manuscript.

To determine whether *aldB* encodes an acetaldehyde dehydrogenase, we first deleted it from the *Z. mobilis* ZM4 genome and observed the effect on growth in oxic and anoxic conditions in rich and minimal media (**Figure 2**). We observed that the deletion had little impact on growth in anoxic conditions but reduced the final culture density by 19% and 30% in oxic rich and minimal media, respectively. We also measured acetate production by both strains under each condition. The wild-type (WT) strain produced acetate in all culture conditions, while acetate production by Δ*aldB* was undetectable except in oxic rich medium (**Figure 3**). In oxic rich medium, Δ*aldB* produced a small amount of acetate (1.37 mM), but this was a 96% decrease compared with the 30.87 mM produced by WT in the same conditions.

**Figure 2.**
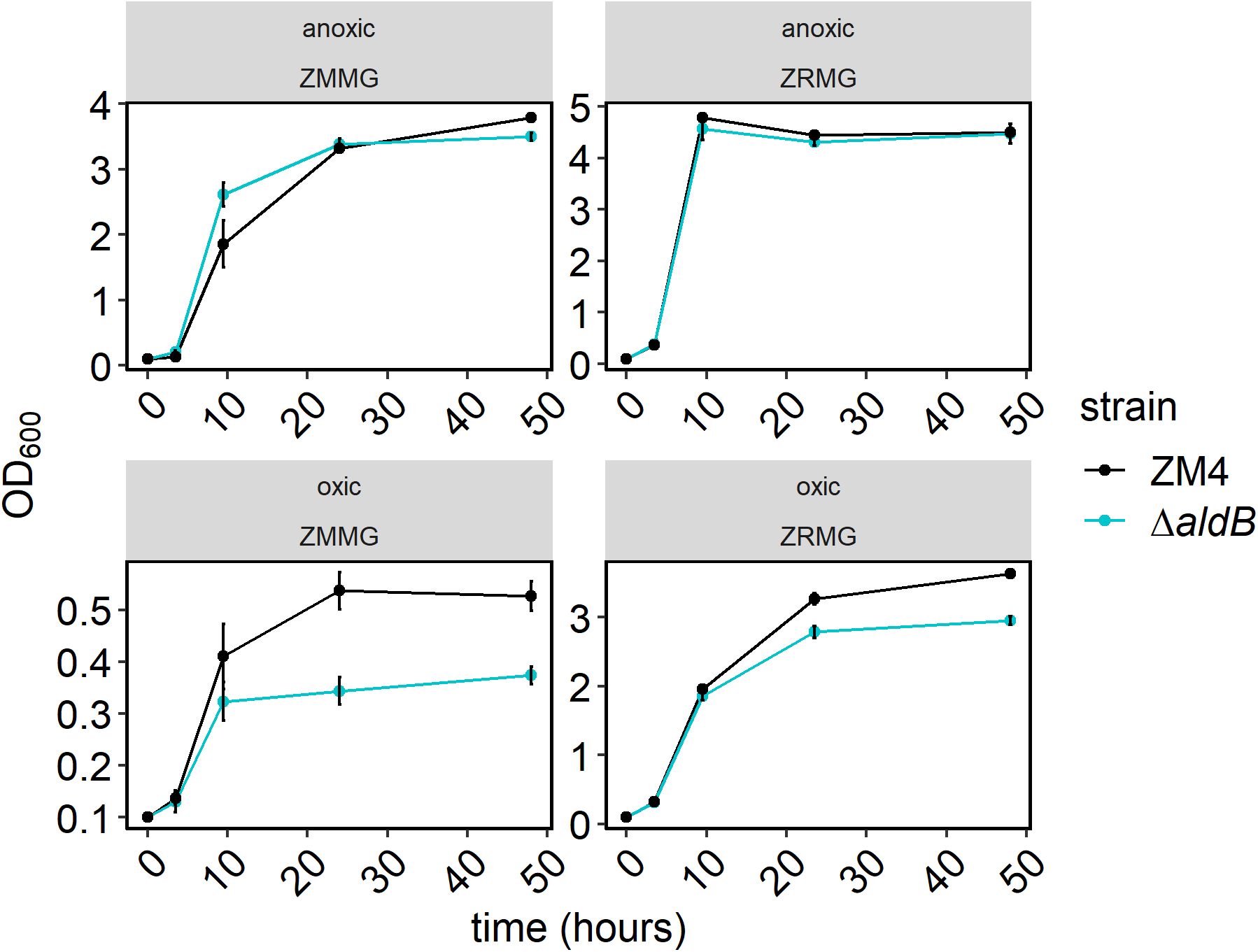
Growth of *Z. mobilis* ZM4 WT and Δ*aldB*. Overnight cultures were inoculated from single colonies and grown overnight in rich or minimal media in static cultures in oxic or anoxic conditions. For subsequent anoxic growth, the cultures were diluted to OD_600_ of 0.1 in 5 ml of fresh rich or minimal media in Hungate tubes in an anaerobic chamber and tubes were closed with stoppers and secured with crimps. For oxic growth cultures were started similarly but Hungate tubes were covered with aluminum foil. All cultures were grown outside anaerobic chamber in 30°C with shaking at 250 rpm for time indicated. Samples were taken periodically to monitor growth. Points represent the average of 3 biological replicates and error bars represent standard error. Note that y-axis scales differ across panels.

**Figure 3.**
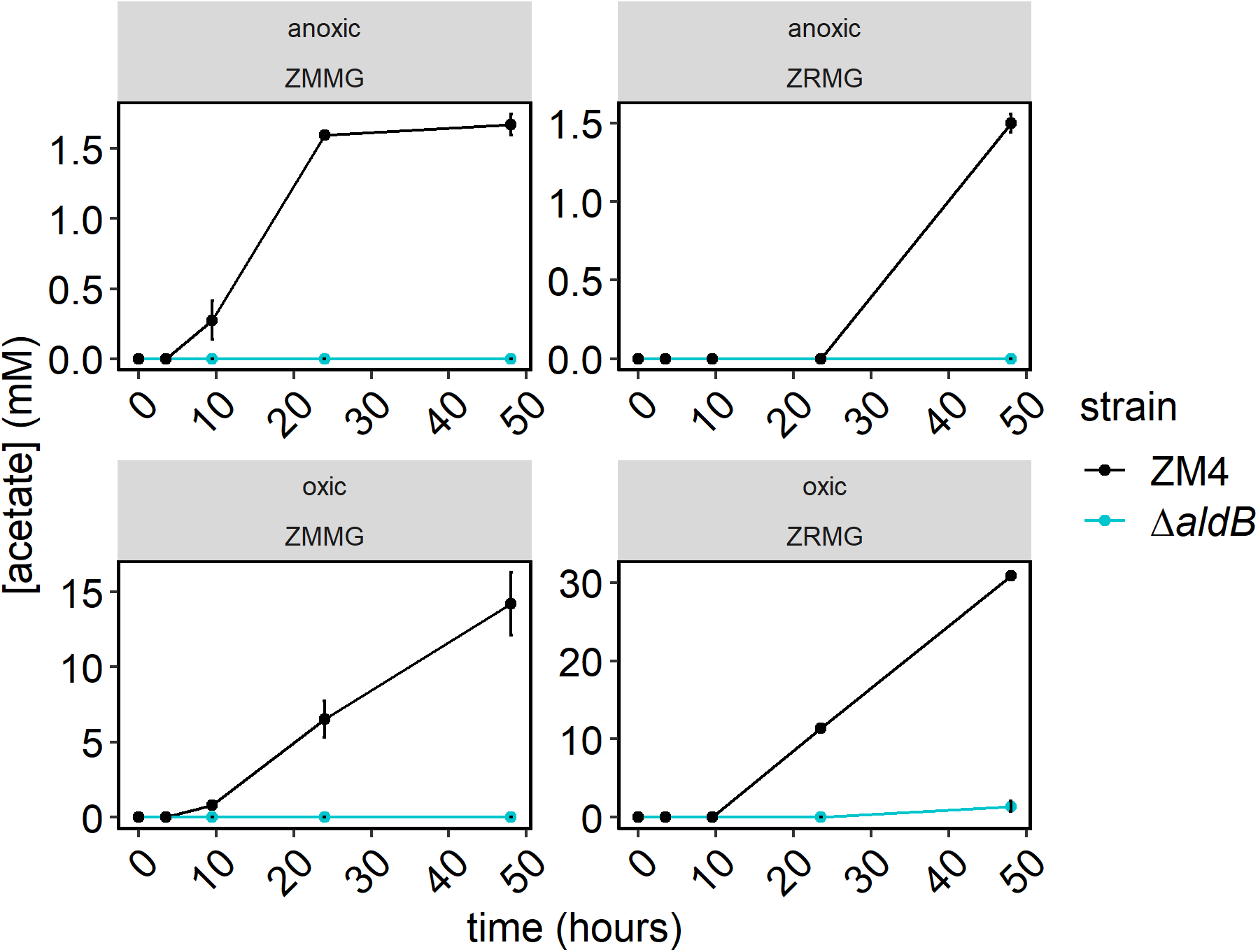
Acetate production by *Z. mobilis* ZM4 WT and Δ*aldB*. Samples from cultures used in Figure 1 were centrifuged and clear supernatants were analyzed by HPLC as described in Materials and Methods. Acetate concentrations were calculated from standard curves. Points represent the average of 3 biological replicates and error bars represent standard error. Note that y-axis scales differ across panels.

We complemented Δ*aldB* by expressing *aldB* from an IPTG inducible plasmid (pRL*aldB*) and measuring growth and acetate production in oxic rich medium (**Figure 4**). We used pRL814, which expresses GFP under an IPTG-inducible promoter, as a control for the effect of the plasmid and heterologous protein expression. We observed that expressing GFP slightly reduced growth in both WT and Δ*aldB*, while expressing *aldB* dramatically improved growth for both strains. Similarly, GFP expression had no effect or reduced acetate production, while *aldB* expression increased acetate production by nearly 3-fold for ZM4 and over 16-fold for Δ*aldB*. Overall, we found that expression of *aldB in trans* complemented the growth and acetate production phenotypes of Δ*aldB*. Further, expression *in trans* increased growth and acetate production in WT under the conditions tested. We also performed complementation with His-tagged AldB and observed the same results, indicating that the tagged version of the protein is active (Figure S4).

**Figure 4.**
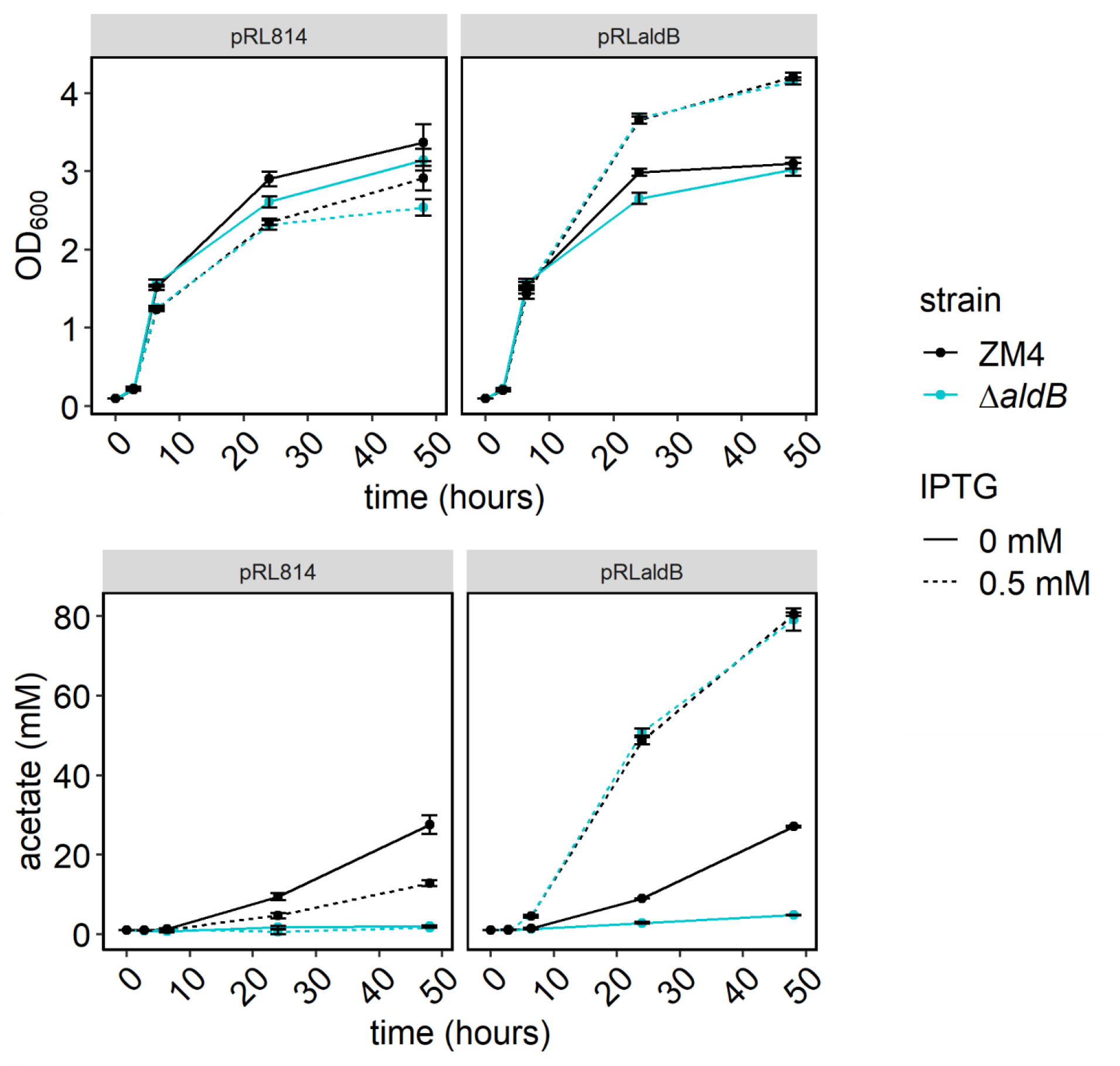
Growth and acetate production by *Z. mobilis* ZM4 WT and Δ*aldB* expressing *aldB* from inducible plasmid. Strains bearing pRL*aldB* or pRL814, were grown as in Figure 1 but media were supplemented with 100 μg/ml of spectinomycin and IPTG was added to 0.5 mM, at time of dilution, when indicated. HPLC analysis was performed as in Figure 2. Points represent the average of three biological replicates and error bars represent standard error.

We confirmed that the Δ*aldB* phenotype was caused by loss of acetaldehyde dehydrogenase activity by using an activity assay to compare soluble protein fractions of WT and Δ*aldB*. We tested activity of the soluble protein fraction to convert acetaldehyde to acetate using NADP^+^ or NAD^+^ as the cofactor. We observed that deletion of *aldB* completely abolished the NADP^+^-dependent activity. In WT, the NAD^+^-dependent acetaldehyde dehydrogenase activity was <20% as high as the NADP^+^-dependent activity. The NAD^+^-dependent activity was reduced slightly by deletion of *aldB* (**Figure 5**).

**Figure 5.**
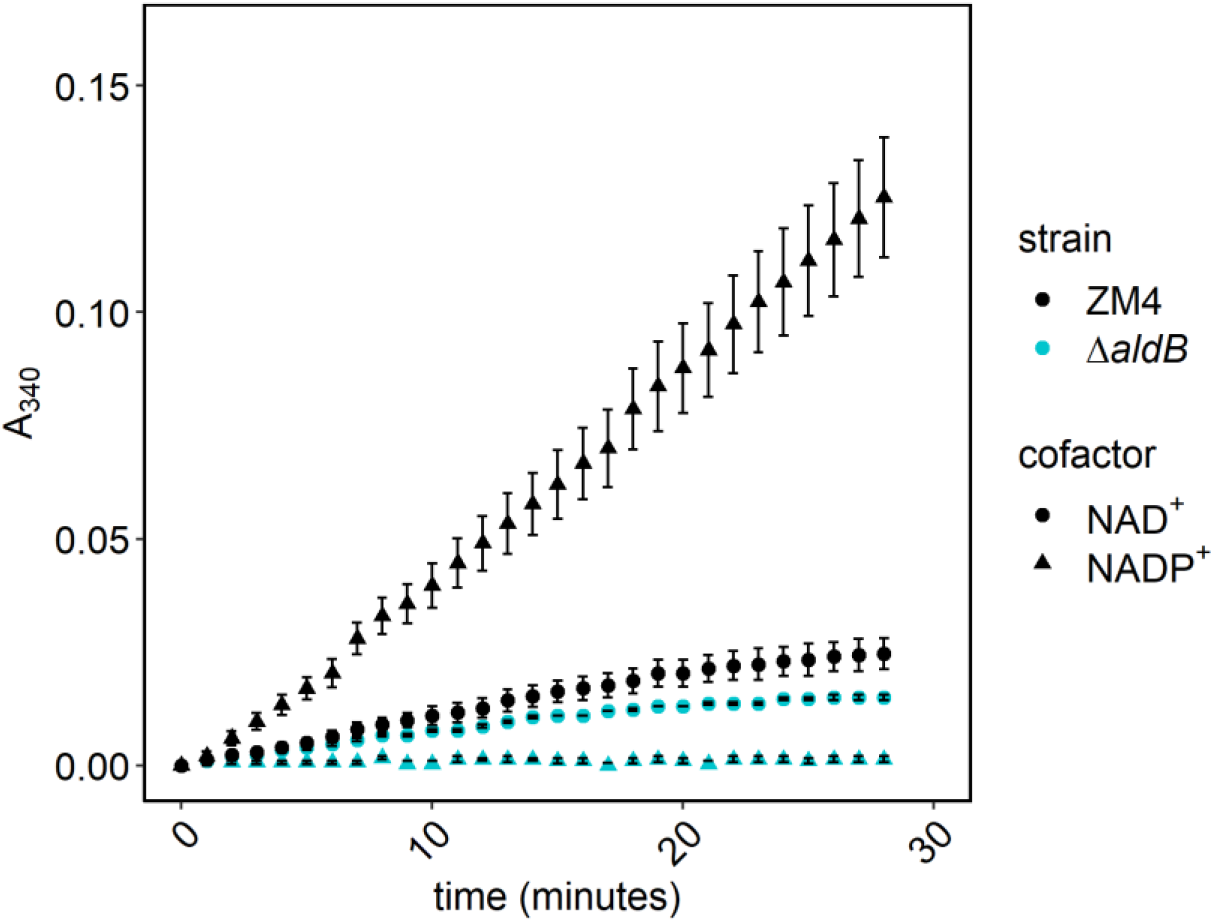
Acetaldehyde dehydrogenase activity in soluble protein fractions from ZM4 WT and Δ*aldB*. Soluble protein fraction (FI) was obtained as described in “Materials and Methods”. Average, total protein concentration in FI was 5.7 +/− 0.5 mg/ml and 5.2 +/− 0.7 mg/ml for ZM4 and Δ*aldB*, respectively. Each enzymatic reaction contained: 0.1 M Tris HCl pH 8.0, 0.1 M KCl, 10 mM β-mercaptoethanol, 2 mM acetaldehyde and 0.67 mM NAD or NADP (protocol for yeast acetaldehyde dehydrogenase from Sigma-Aldrich). Reaction was started by adding 33 μl of FI and measured for 30 minutes at 25°C in 24-well microtiter plate as described in “Materials and Methods”. Absorbance at 340 nm in control without FI was subtracted from the reactions. Points represent the average of three independent experiments with three technical repeats and error bars represent standard error.

We next overexpressed a His-tagged version of AldB in *E. coli* and confirmed acetaldehyde dehydrogenase activity of the tagged protein in the soluble protein fraction from cells induced with IPTG (**Figure 6**). We purified the protein using Ni-affinity chromatography as described in Materials and Methods (Figure S3). We pooled fractions based on their specific activity and used fractions 9-11 to measure substrate specificity using an acetaldehyde dehydrogenase assay as described in Materials and Methods. We observed that the activity was specific for acetaldehyde with moderate activity toward other short-chain fatty aldehydes, propionaldehyde (3C), butyraldehyde (4C), and valeraldehyde (5C) (**Table 3**). The activity toward formaldehyde and glyceraldehyde was low.

**Table 3.**
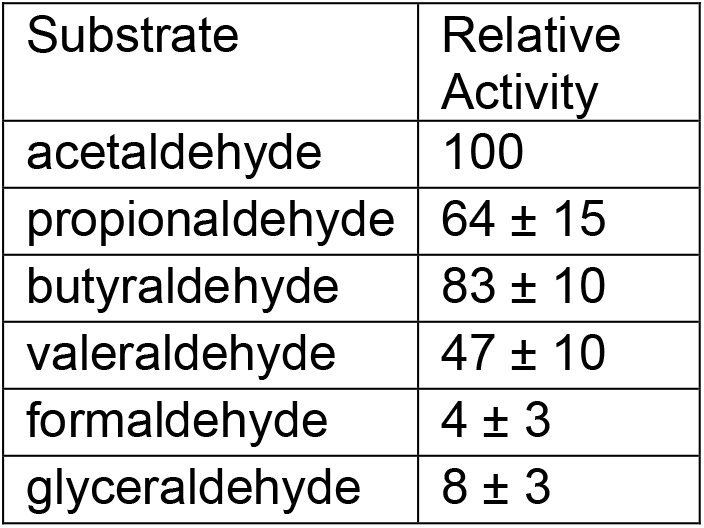
Substrate specificity of ZM4 acetaldehyde dehydrogenase. Enzymatic reactions were performed in standard assay conditions (22 mM aldehyde, 2 mM NADP^+^, 10 mM DTT, 50 mM Tris HCl pH 8.0 and 50 mU of enzyme) at 30°C as described in Materials and Methods for 10 minutes. Aldehydes were added as 0.23 M solutions in DMSO. Activity was normalized by setting activity with acetaldehyde at 100%. Data are averages of 5 independent experiments with three technical replicates for each substrate.

| Substrate | Relative Activity |
| --- | --- |
| acetaldehyde | 100 |
| propionaldehyde | 64 ± 15 |
| butyraldehyde | 83 ± 10 |
| valeraldehyde | 47 ± 10 |
| formaldehyde | 4 ± 3 |
| glyceraldehyde | 8 ± 3 |

**Figure 6.**
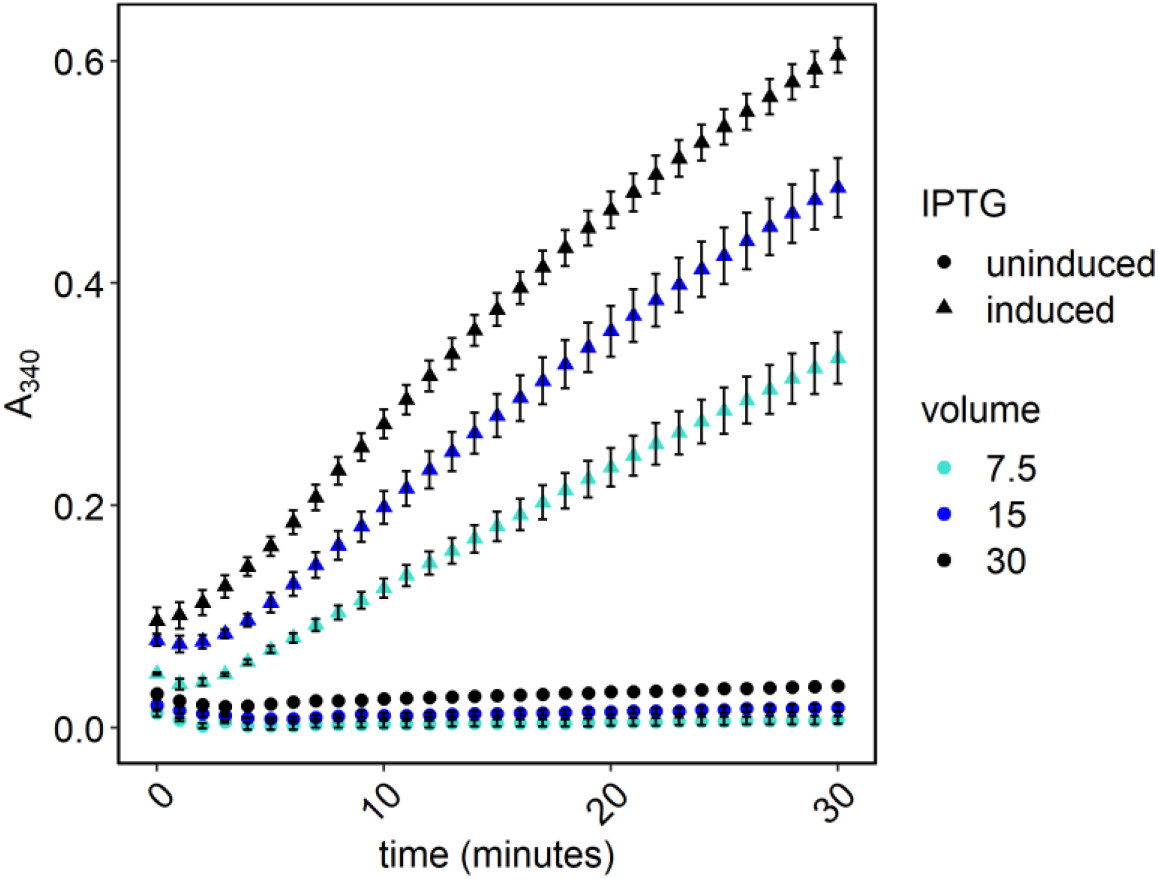
Acetaldehyde dehydrogenase activity in soluble protein fraction from *E. coli* BL21 carrying a plasmid expressing *aldB*. 50-mL cultures were grown in LB supplemented with ampicillin at 30°C to OD_600_ 0.4. At this point, IPTG was added to a final concentration of 0.1 mM, where indicated, and growth continued for 1.5 hours. FI was obtained as described in “Materials and Methods” for *Z. mobilis*. Average total protein concentration in three independent FI was 11±1.2 mg/ml and 8.4±1.1 mg/ml for uninduced and induced cultures, respectively. An enzymatic assay was performed as described in Figure 4, with NADP as a cofactor. The reaction was started by adding variable volumes of FI as indicated. Points represent the average of three independent FI and error bars represent standard error.

Because we previously observed some acetate secretion by ZM4 grown in lignocellulosic hydrolysate, we also measured growth and acetate concentration in fermentations using ZM4 or Δ*aldB*. Growth and substrate utilization by the two strains was not significantly different; both strains reached a final OD_600_ of 5.4 after 48 hours and consumed all available glucose. However, the acetate and ethanol concentrations were significantly different between the two strains. The initial acetate concentration in the hydrolysate was 38.48 mM and after 48 hours of fermentation by ZM4, the concentration rose to 41.31±0.06 mM. In contrast, with Δ*aldB*, the acetate concentration decreased slightly to 37.76±0.04 mM. Ethanol production by Δ*aldB* was slightly decreased, at 750.01±0.28 mM versus 752.7±0.21 mM for ZM4.

## Discussion

We determined that the locus ZMO1754 (also known as ZMO_RS07890) in the *Z. mobilis* ZM4 genome encodes an acetaldehyde dehydrogenase that is the main source of acetate production. Here, we refer to the gene encoded at that locus as *aldB* based on functional similarity to *E. coli aldB*. Although acetate production by *Z. mobilis* is well-documented and an acetaldehyde dehydrogenase was previously purified from strain Z6, the genes necessary for acetate production were previously unknown. Yang et al. previously speculated that ZMO1754 could encode an acetaldehyde dehydrogenase but did not test the hypothesis because they found no evidence of upregulation in response to oxygen (5). However, a more recent transcriptomic and proteomic characterization of oxygen response in *Z. mobilis* did observe that the transcript and protein encoded by ZMO1754 were upregulated in response to oxygen (18). The newer multi-omic characterization led us to hypothesize that ZMO1754 encodes an acetaldehyde dehydrogenase. Our deletion studies show that ZMO1754 encodes the enzyme responsible for acetate production in *Z. mobilis* ZM4.

We also characterized AldB from *Z. mobilis* ZM4 to determine whether it is an ortholog of the NADP^+^-dependent acetaldehyde dehydrogenase purified from *Z. mobilis* Z6 by Barthel et al. (11). We found that both are aldehyde dehydrogenases with significant specificity toward acetaldehyde, are upregulated in response to oxygen, have approximately the same molecular weight, and use NADP^+^ rather than NAD^+^ as a cofactor. Although we observed a minor NAD^+^-dependent acetaldehyde dehydrogenase activity in the protein fraction extracted from Δ*aldB*, both acetate concentrations in culture and activity assays confirm that *aldB* encodes the major acetaldehyde dehydrogenase.

The presence of a growth defect in Δ*aldB* only under oxic conditions suggests that acetate production is beneficial only when oxygen is present. This finding aligns with previous research indicating that acetaldehyde accumulation causes poor growth of *Z. mobilis* when oxygen is present (**Figure 1**) (20, 21). We observed that although there was some acetate production under anoxic conditions, the concentration was >10-fold lower than in oxic conditions, suggesting that there is little acetaldehyde available as a substrate when oxygen is absent. In context with previous work, our findings suggest that the main role of AldB is to detoxify acetaldehyde that is generated under oxic conditions.

From a biotechnological perspective, identifying the locus encoding acetaldehyde dehydrogenase is useful because a deletion strain could be used to eliminate acetate production in biofuel fermentations. On the other hand, we observed decreased ethanol production by Δ*aldB*, suggesting that acetate production may be involved in a stress-response pathway in *Z. mobilis* ZM4. We speculate that reducing equivalents may be diverted from central metabolism to detoxification pathways that process toxins found in lignocellulosic hydrolysates. The diversion of reducing equivalents would necessitate acetaldehyde detoxification as observed in oxic conditions. Further, *Z. mobilis* has been proposed as a host for acetaldehyde production and deletion of *aldB* is likely to dramatically increase acetaldehyde yield (22). (Note that we do not report acetaldehyde concentrations here because our culture method resulted in significant evaporative loss of acetaldehyde.) Further work is necessary to determine whether deletion or upregulation of *aldB* from *Z. mobilis* would be beneficial for biotechnological processes overall.

We expect our findings to be generalizable across *Zymomonas* because the acetaldehyde dehydrogenase encoded by *aldB* in strain ZM4 appears to be conserved across the entire *Zymomonas* genus. We recently conducted a genome comparison analysis for all sequenced *Zymomonas* genomes and used our previous ortholog analysis to determine that an *aldB* ortholog is present in all analyzed genomes. The *aldB* gene shares 99.1% identity across *Z. mobilis* subspecies *mobilis* and 92% and 84% between *francensis* and *mobilis* and *pomaceae* and *mobilis*, respectively (7).

## Acknowledgements

We thank Dylan Courtney, Dr. Jennifer Reed, Dr. Brian Pfleger, and Dr. Yaoping Zhang, (University of Wisconsin, Madison) for helpful discussions and Nicholas Tefft for help with HPLC analysis. This material is based upon work supported by the Great Lakes Bioenergy Research Center, U.S. Department of Energy, Office of Science, Office of Biological and Environmental Research under Award Number DE-SC0018409.

